# SUMO mediates the coordinate regulation of meiotic chromosome length and crossover rate

**DOI:** 10.64898/2026.03.10.710713

**Authors:** Yan Yun, Huanyu Qiao, Martin White, Sumit Sandhu, Wendy Qiu, Sarah Bourne, Anusha Deshpande, Shubhang Bhatt, Ajay Sharma, Logan Bailey, Hung Tran, Ben Van, H.B.D. Prasada Rao, Neil Hunter

## Abstract

Meiotic prophase-I chromosomes are organized into linear arrays of chromatin loops anchored to proteinaceous axes that define the interaction interfaces for the pairing and synapsis of homologous chromosomes. Chromatin loop size and axial chromosome length are inversely correlated and vary widely both between and within species, including between the sexes. The molecular basis of this variation remains unclear. Here, we provide evidence that the small ubiquitin-like modifier, SUMO, regulates loop–axis organization in mouse meiosis. Our analysis shows that the longer axes of oocyte chromosomes contain more SUMO per unit length than the shorter axes of spermatocyte chromosomes. In mouse models, the loss of SUMO1 results in shorter axes and longer chromatin loops. Conversely, increased SUMO1 conjugation, caused by mutation of the SENP1 isopeptidase, produces longer axes with shorter loops. Axis length positively correlates with meiotic recombination. Accordingly, *Sumo1* and *Senp1* mutations respectively decrease and increase crossover frequency. These findings identify SUMO as a key regulator of meiotic chromosome architecture and suggest a molecular basis for the physiological variation in chromosome length and recombination rates seen among species, sexes, individuals, and individual meiocytes.

**GRAPHICAL ABSTRACT:** 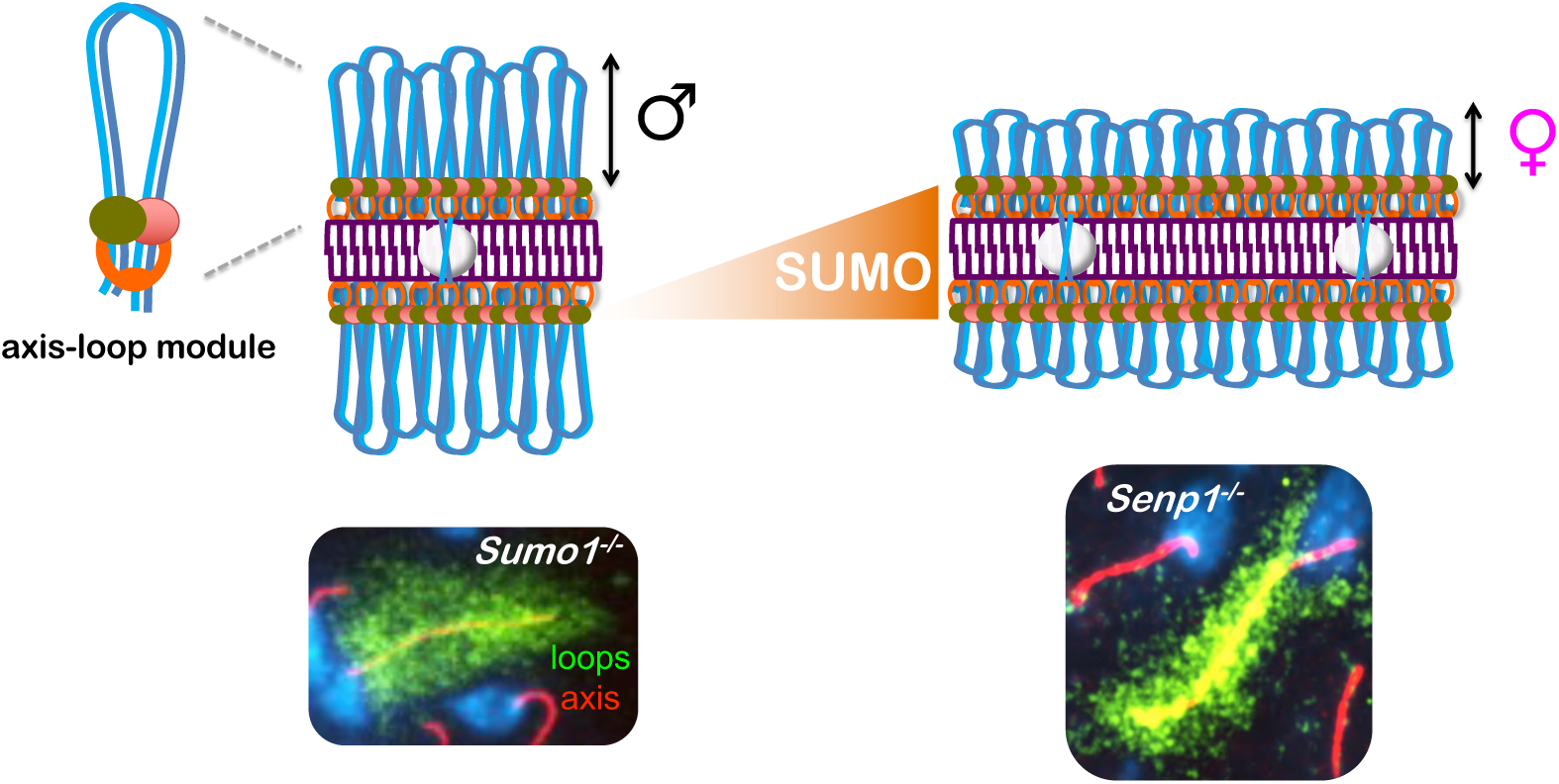

## INTRODUCTION

During meiotic prophase I, the conjoined sister-chromatids of each replicated chromosome (known as a homolog) organize into dual linear arrays of co-orientated chromatin loops, the bases of which coalesce into a structural axis that defines the interface for pairing, synapsis, and recombination with a homologous partner chromosome (1–3). The physical association and functional interplay between homolog axes and recombination factors govern each step of meiotic recombination including the formation of DNA double-strand breaks (DSBs), inter-homolog template bias, and the choice between crossover and non-crossover outcomes (4).

A striking manifestation of this interplay is the positive correlation between the axial lengths of prophase-I chromosomes and recombination rates (5, 6). Covariation of axis length and recombination rate is seen in several physiological contexts in which the genome size is fixed; for example between closely related species (7, 8), different strains of the same species (9, 10), males and females of the same species (9, 11–13), individuals of the same sex (9, 14), and even between different meiocytes from a single individual (6, 14). These observations suggest that heritable, gonad-specific (e.g., ovary vs. testis), and cell-autonomous factors can all contribute to the covariation of axis length and recombination rate. How axes length is regulated remains unclear. Here we reveal a role for the small ubiquitin-like modifier, SUMO, in regulating chromosome axis length during meiosis in mouse.

## RESULTS

### Sexually Dimorphic Localization of SUMO Along Prophase-I Chromosomes

Crossover rate is sexually dimorphic in many species (also known as heterochiasmy), including human and mouse, and positively correlates with global differences in the axial lengths of synapsed meiotic prophase-I chromosomes (5, 6, 9, 11–13, 15, 16). This relationship was corroborated in mouse by immunofluorescence analysis of surface-spread prophase-I chromosomes from spermatocytes and oocytes (**Figure 1A–C**). Immunostaining for the homolog axis component SYCP3 and the crossover marker MLH1 indicated that pachytene-stage oocytes chromosomes were on average 22.2% longer (total axis length of 253.8 ± 23.5 µm versus 207.7 ± 31.9 µm, mean ± S.D.; *p*<0.0001, Mann-Whitney test) and had 16.6% more crossovers than spermatocyte chromosomes at the same stage (26.0 ± 1.5 versus 22.3 ± 1.5 in; *p*<0.0001, unpaired *t* test).

**Figure 1.**
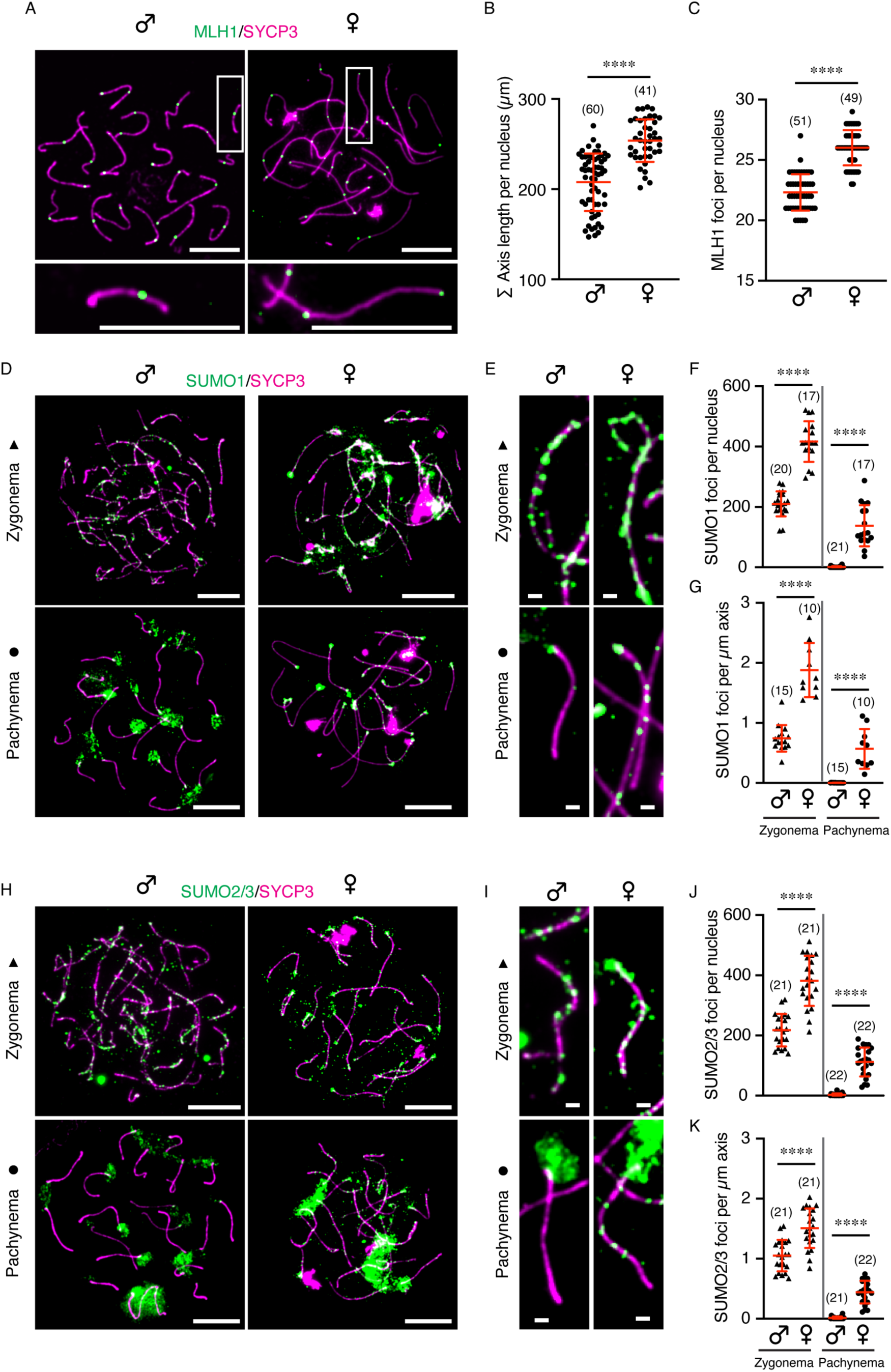
Sexually dimorphic localization of SUMO along prophase-I chromosomes. **A**. Representative images showing chromosome axis marker SYCP3 (magenta) and crossover marker MLH1 (green) in mouse pachytene spermatocytes and oocytes, respectively. Chromosomes in white rectangle are enlarged in the bottom panels. Scale bars are 10 μm (top) and 1 μm (bottom). **B, C**. Quantification of total axis length (B) or MLH1 foci (C) per spermatocyte and oocyte. **D, H**. Representative images of SUMO1 (D) or SUMO2/3 (H) in zygotene and pachytene nuclei of both sexes. Scale bars are 10 μm. **E, I**. Representative homologous chromosome axis showing SUMO1 (E) or SUMO2/3 (I) foci in zygotene and pachytene nuclei of spermatocyte and oocyte, respectively. Scale bars are 1 μm. **F, J**. Quantification of SUMO1 (F) or SUMO2/3 (J) foci along chromosome axis per nucleus. **G, K**. Comparison of SUMO1 (G) or SUMO2/3 (K) foci density along chromosome axis between spermatocyte and oocyte. Data are presented as mean ± SD (B, C, F, G, J, K). Data were analyzed with Mann-Whitney test or unpaired t test. Numbers in parenthesis indicate number of nuclei analyzed. \*\*\*\**p* < 0.0001.

We previously demonstrated roles for the SUMO-modification system in regulating meiotic recombination in mouse and showed that SUMO localizes along synapsing chromosomes in spermatocytes (17). Immunostaining for the SUMO1 isoform revealed 210.3 ± 41.6 foci per nucleus along the axes of zygotene-stage spermatocytes but foci almost completely disappeared by early pachytene (1.0 ± 2.5, mean ± S.D.; **Figure 1D–F**). At the same time, SUMO1 accumulated into large staining structures localized at centromeric heterochromatin. Strikingly, two-fold more SUMO1 foci were detected along the chromosome axes of zygotene-stage oocytes (416.6 ± 67.2 per nucleus, mean ± S.D.; *p*<0.0001 relative to zygotene spermatocytes, unpaired *t* test; **Figure 1D–F**). This increase was not simply a result of the longer chromosome axes in oocytes because the density of foci per µm of axis was also much higher (1.9 ± 0.5 in oocytes versus 0.7 ± 0.2 in spermatocytes, mean ± S.D.; *p*<0.0001, unpaired *t* test; **Figure 1G**). Moreover, unlike spermatocytes, SUMO1 foci persisted in pachytene-stage oocytes (*p*<0.0001 for comparisons of both focus numbers and focus densities, Mann-Whitney test; **Figure 1D–F**). Very similar sexually dimorphic localization patterns were seen for the SUMO2/3 isoforms (antibodies do not distinguish these two closely-related isoforms; **Figure 1H–K**). Thus, SUMO is both more abundant and more persistent along the chromosome axes of prophase-I oocytes than those of spermatocytes.

### Chromosome Length During Prophase-I is Modulated by SUMO

Given the positive correlation between chromosome length and SUMO levels, we sought to address whether there is a functional relationship. To this end, we analyzed two mouse lines that alter SUMO1 levels in opposite ways (**Figure 2**). *Sumo1^Gt(XA024)Byg^* mice (hereafter *Sumo1^Gt^*) carry a gene-trap allele, and homozygous animals completely fail to express the SUMO1 protein (18). Absence of SUMO1 in meiotic prophase-I was confirmed by immunostaining oocyte and spermatocytes chromosome spreads from *Sumo1^Gt/Gt^* homozygous animals (**Figure 2A–C** and **Supplemental Figure S1**). Conversely, the *Senp1^M1Mku^* proviral-integration allele (hereafter *Senp1^M1)^* is a severe hypomorph for SENP1, the SUMO1-specific isopeptidase, resulting in greatly elevated levels of SUMO1 conjugates (19, 20). In *Senp1^M1/M1^* homozygotes, SUMO1 immunostaining intensity was 2.6- and 2.7-fold higher than wild-type intensities in zygotene-stage spermatocytes and oocytes, respectively (mean ± S.D., *p*=0.0028 and <0.0001, respectively; **Figure 2A–C**). Moreover, SUMO1 remained hyperabundant during pachytene (**Supplemental Figure S1**).

**Figure 2.**
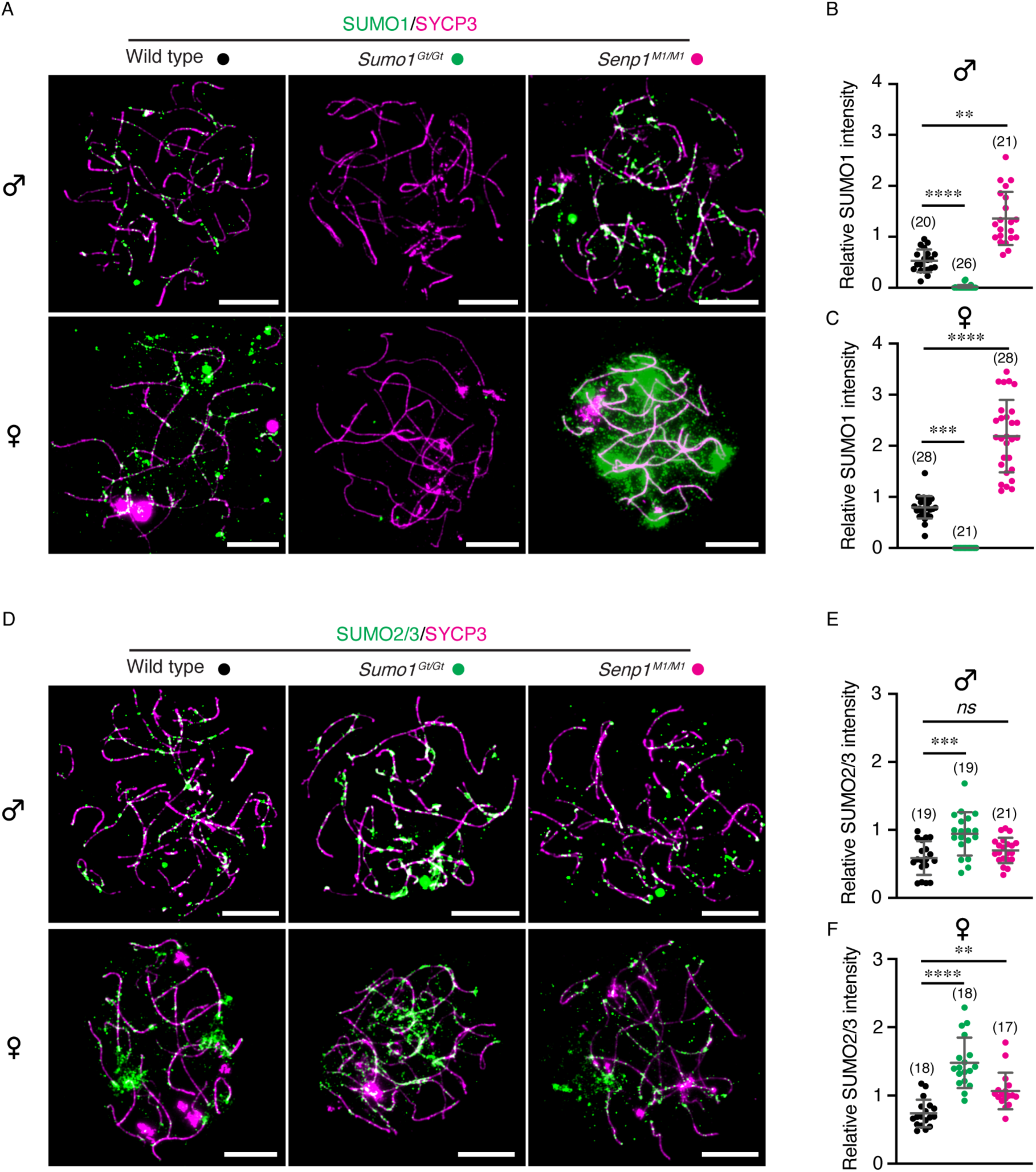
SUMO levels in both spermatocytes and oocytes of gene-modified mouse models. **A, D**. Representative immunostaining images of SUMO1 (A) or SUMO2/3 (D) of zygotene spermatocyte and oocyte nuclei, from wild type, *Sumo1^Gt/Gt^*, and *Senp1^M1/M1^* mice, respectively. Scale bars are 10 μm. **B, C**. Quantification of relative SUMO1 signal per nuclei of spermatocyte (B) or oocyte (C). **E, F**. Quantification of relative SUMO2/3 signal per nuclei of spermatocyte (E) or oocyte (F). Relative SUMO1 and SUMO2/3 intensity were calculated as signal intensity ratios of total axis-associated signal for SUMO divided by the signal for SYCP3. Data are presented as mean ± SD (B, C, E, F). Data were analyzed with Kruskal-Wallis test followed by Dunn’s multiple comparisons (B, C, F), and ordinary one-way ANOVA followed by Dunnett’s multiple comparisons (E). Numbers in parenthesis indicate number of nuclei analyzed. ns, not significant; \*\**p* < 0.01; \*\*\**p* < 0.001; \*\*\*\**p* < 0.0001.

SUMO2/3 are inferred to substitute for SUMO1 in *Sumo1^Gt/Gt^*animals, largely compensating for its absence and thereby preventing overt phenotypes (18). Also, in mixed oligomers, SUMO1 is thought to cap SUMO2/3 chains and thereby limit their lengths (20, 21). Consistent with both inferences, SUMO2/3 staining intensity was increased in *Sumo1^Gt/Gt^* meiocytes (1.6- and 2.0-fold above wild-type intensities in zygotene-stage spermatocytes and oocytes, respectively; mean ± S.D., *p*=0.0001 and <0.0001, respectively; **Figure 2D–F**). SUMO2/3 intensities were also slightly elevated in *Senp1^M1/M1^*meiocytes (*p*=0.0096; **Figure 2D–F**). This observation could reflect a greater stability of SUMO2/3 chains that are capped by SUMO1 (20).

Synapsis defects were not detected in either *Sumo1^Gt/Gt^*or *Senp1^M1/M1^* meiocytes. However, measurements of chromosome axis lengths revealed a striking relationship with SUMO1 (**Figure 3A–C**). Absence of SUMO1 in *Sumo1^Gt/Gt^* meiocytes resulted in shorter chromosomes in both males and females (189.4 ± 31.3 µm versus 207.7 ± 31.9 µm in wild-type spermatocytes, mean ± S.D., *p*=0.0076, **Figure 3B**; and 217.1 ± 21.9 µm versus 272.4 ± 30.5 µm in wild-type oocytes, mean ± S.D., *p*<0.0001, **Figure 3C**). The magnitude of chromosome shortening was greater in oocytes (20.3%) than in spermatocytes (8.8%).

**Figure 3.**
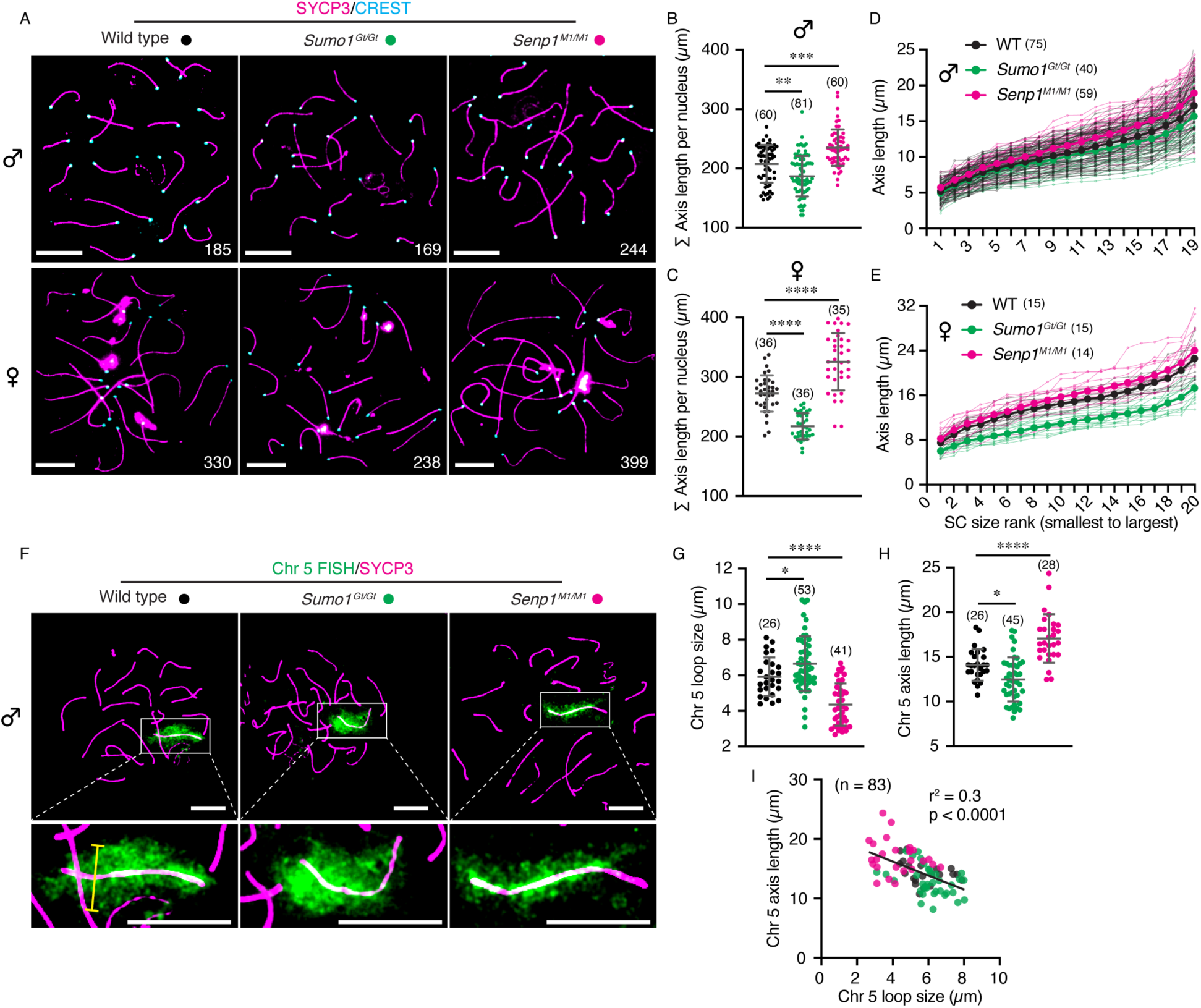
Chromosome organization during prophase-I is modulated by SUMO1 Levels in both sexes. **A**. Representative immunostaining images from surface spreads of pachytene spermatocytes and oocytes, in wild type, *Sumo1^Gt/Gt^*, and *Senp1^M1/M1^* mice, respectively. White numbers are total axis length per nucleus. Scale bars are 10 μm. **B, C**. Quantification of total homologous chromosome axis length measured in pachytene spermatocytes (B; 19 autosome chromosomes) and oocytes (C; 20 chromosomes) from different mouse models. **D, E**. Line plots of axis length versus individual homologous chromosome (from small to large size) in a pachytene spermatocyte (D) or oocyte (E) from different mouse models. Bold lines indicate average value of the corresponding chromosomes from a specific genotype. **F**. Representative images of pachytene spermatocyte chromosome spreads labelled with FISH signal of whole chromosome 5, from different mouse models. Labelled chromosomes in white rectangles are enlarged in the bottom panels. Scale bars are 10 μm (top and bottom). Yellow line indicates measurement of loop size. **G, H**. Quantification of chromosome 5 loop size (G) or its axis length (H) in pachytene spermatocytes of different mouse models. Loop size is calculated as an average value of 4-11 measurements for each chromosome depending on the axis length. **I**. Negative association between chromosome 5 axis length and its loop size in pachytene spermatocytes. Simple linear regression, r^2^ = 0.3; *p* < 0.0001. Data are presented as mean ± SD (B, C, G, H). Data were analyzed with Kruskal-Wallis test followed by Dunn’s multiple comparisons (B), Brown-Forsythe and Welch ANOVA followed by Dunnett’s T3 multiple comparisons (C), and ordinary one-way ANOVA followed by Dunnett’s multiple comparisons (G, H). Numbers in parenthesis indicate number of nuclei analyzed. \**p* < 0.05; \*\**p* < 0.01; \*\*\**p* < 0.001; \*\*\*\**p* < 0.0001.

Oppositely, excess SUMO1 conjugation in the *Senp1^M1/M1^* mutant was associated with longer chromosome axes (235.2 ± 30.8 µm versus 207.7 ± 31.9 µm in wild-type spermatocytes, mean ± S.D., *p*=0.0004, **Figure 3B**; and 325.7 ± 48.2 µm versus 272.4 ± 30.5 µm in wild-type oocytes, mean ± S.D., *p*<0.0001, **Figure 3C**). Again, the effect was greater in oocytes (19.6% length increase) than spermatocytes (13.2%).

Analysis of individual chromosome lengths indicated that the effects of *Sumo1^Gt/Gt^* and *Senp1^M1/M1^* mutations were global, i.e. length changes were not confined to one or a few chromosomes (**Figure 3D,E**). Although axis lengths were measured in mid-late pachytene nuclei containing mature crossover sites marked by MLH1 foci, length differences in *Sumo1^Gt/Gt^*and *Senp1^M1/M1^* mutants were already present in early pachytene nuclei (**Supplemental Figure S2**), possibly reflecting early perturbations of meiotic chromosome organization.

Variation in the axial lengths of meiotic prophase-I chromosomes can occur through changes in chromatin organization, specifically the sizes of chromatin loops. Given that the density of chromatin loops along prophase-I homolog axes is evolutionarily conserved (at ∼25 loops per µm of axis)(22), changes in axis length show an inverse relationship with the lengths of chromatin loops, in both physiological and mutant situations (3, 5, 11, 12, 23–28). To estimate chromatin loop lengths in *Sumo1^Gt/Gt^* and *Senp1^M1/M1^* mutant spermatocytes, a whole-chromosome FISH probe was used to visualize mouse chromosome 5 and the widths of the FISH-signals were measured (**Figure 3F,G**). Co-staining for SYCP3 allowed the axial lengths of the same chromosomes to be measured (**Figure 3H**). The shorter chromosomes observed in *Sumo1^Gt/Gt^* spermatocytes correlated with 12.3% wider FISH signals implying longer chromatin loops (6.7 ± 1.5 µm in *Sumo1^Gt/Gt^* versus 5.9 ± 1.1 µm in wild type, mean ± S.D.; *p*=0.0444; **Figure 3G**). Conversely, in *Senp1^M1/M1^* spermatocytes, FISH signals were 26.4% narrower (4.4 ± 1.2 µm in *Senp1^M1/M1^* versus 5.9 ± 1.1 µm in wild type, mean ± S.D.; *p*<0.0001; **Figure 3G**). Moreover, a clear negative correlation between axis length and loop size was seen for individual chromosomes (**Figure 3I**).

We infer that increased SUMO1 conjugation results in smaller chromatin loops and thus longer meiotic prophase-I chromosomes. One prediction of this inference is that the longer chromosomes in *Senp1^M1M1^* meiocytes should be SUMO1 dependent. This prediction was tested by measuring chromosome lengths in spermatocytes from *Sumo1^Gt/Gt^ Senp1^M1/M1^* double mutants (**Supplemental Figure S3A**), which confirmed that the longer axis of *Senp1^M1/M1^* spermatocyte chromosomes were largely, though not completely, dependent on the presence of SUMO1. Total axis lengths in *Sumo1^Gt/Gt^ Senp1^M1/M1^* double mutant nuclei averaged 212.6 ± 36.7 µm, significantly smaller than in the *Senp1^M1/M1^* single mutant (235.2 ± 30.8 µm, mean ± S.D.; *p*=0.0044), and very similar to the lengths of wild-type chromosomes (207.7 ± 31.9 µm, mean ± S.D.; *p*>0.9999), but still longer than in *Sumo1^Gt/Gt^* single-mutant nuclei (189.4 ± 31.3 µm, mean ± S.D.; *p*=0.0215; **Supplemental Figure S3A**). Thus, *Senp1^M1^*mutation appears to affect prophase-I chromosome length through SUMO1-dependent and independent pathways.

### Crossover Rate is Modulated by SUMO

In wild-type mammals, including humans, crossover rates and prophase-I chromosome lengths are positively correlated in comparisons between the sexes, individuals of the same sex, and even between individual nuclei (6, 9, 12, 29). Therefore, we tested whether the chromosome-length changes observed in *Sumo1^Gt/Gt^* and *Senp1^M1/M1^* mice were associated with altered rates of crossing over (**Figure 4**). Crossover sites in prophase-I chromosome spreads were detected by immunostaining pachytene-stage nuclei for the crossover-specific marker MLH1 (**Figure 4A**; results were confirmed using additional crossover-specific markers, HEI10 and CDK2, **Supplemental Figure S4**). In *Sumo1^Gt/Gt^* meiocytes, crossover numbers were reduced relative to wild type, from 22.3 ± 1.5 to 20.2 ± 3.4 MLH1 foci in spermatocytes (9.4% reduction); and from 25.8 ± 1.8 to 23.7 ± 1.5 in oocytes (8.1% reduction; mean ± S.D., *p*=0.0019 and <0.0001, respectively; **Figure 4A,B** and **4A,D**). Analysis of crossover numbers per chromosome pair (MLH1 foci per synaptonemal complex, SC) revealed significantly altered distributions in *Sumo1^Gt/Gt^* nuclei (*p*<0.0001 for both spermatocytes and oocytes, Fisher’s exact test, **Figure 4C** and **4E**). In spermatocytes, SCs with two MLH1 foci were reduced from 22.0% in wild type to 19.1% in *Sumo1^Gt/Gt^*, while apparent non-exchange SCs with zero MLH1 foci increased from 2.9% to 8.2% (**Figure 4C**). In oocytes, SCs with three MLH1 foci are occasionally observed in wild type (4.4%) but were absent in *Sumo1^Gt/Gt^* nuclei; SCs with two foci were reduced from 46.5% to 31.0%; while SCs with a single focus increased from 48.8% to 65.5% (**Figure 4E**). SCs lacking MLH1 foci were rarely observed in wild-type oocytes (0.3%) but represented 3.5% of SCs in *Sumo1^Gt/Gt^*.

**Figure 4.**
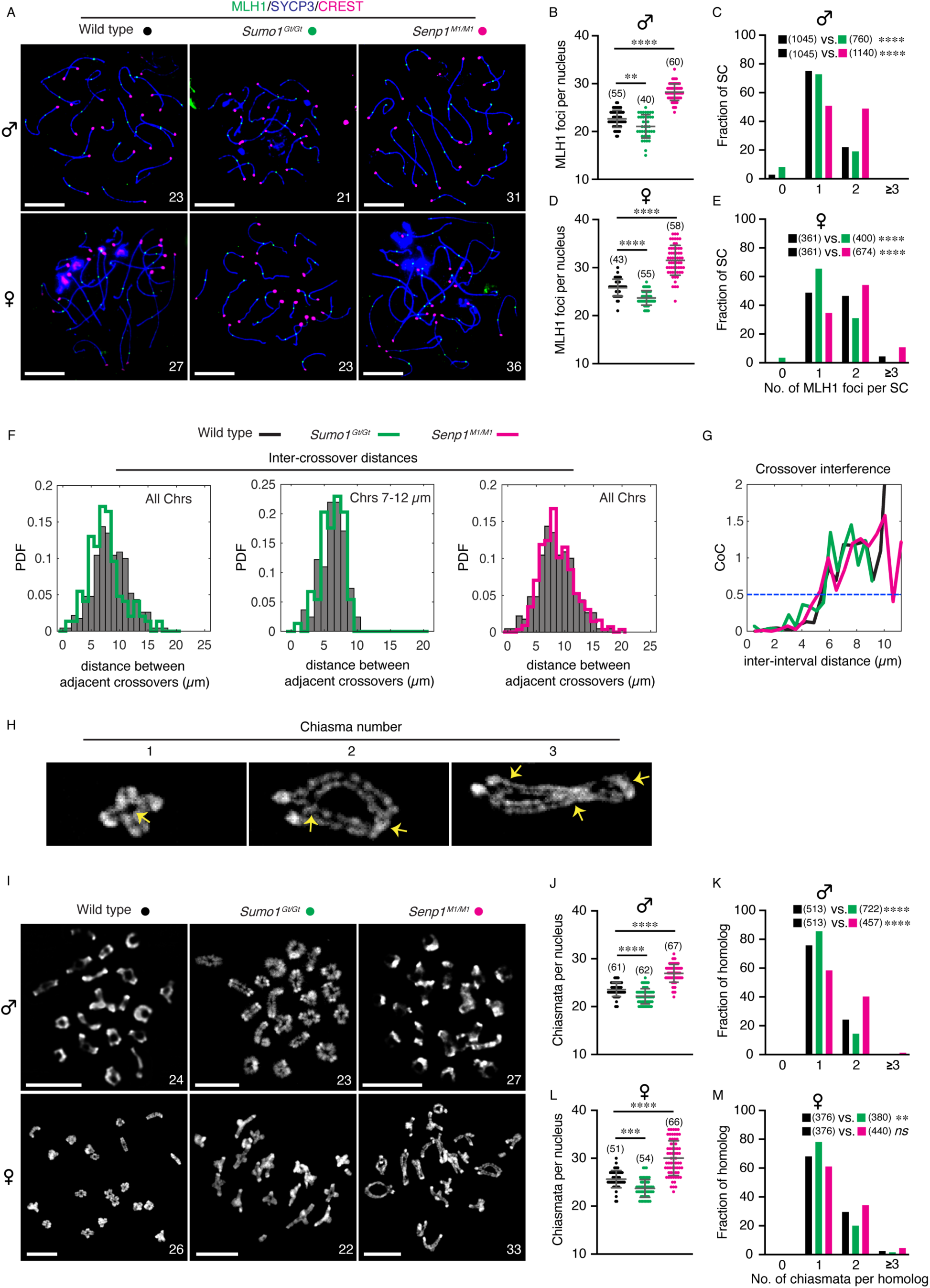
Crossover rate in both spermatocytes and oocytes is regulated by SUMO1 levels. **A**. Representative surface spread images of pachytene spermatocytes and oocytes immunostained for SYCP3 (blue), CREST (magenta), and MLH1 (green) from wild type, *Sumo1^Gt/Gt^*, and *Senp1^M1/M1^* mice, respectively. White numbers indicate MLH1 foci counts per nucleus (19 autosomes in spermatocytes and all 20 chromosomes in oocytes). Scale bars are 10 μm. **B, D**. Quantification of MLH1 foci counts per nucleus of pachytene spermatocytes (B) or oocytes (D) from different mouse genotypes. **C, E**. Fraction of individual homologous chromosomes possessing 0, 1, 2, or more than 3 MLH1 foci, from pachytene spermatocytes (C) or oocytes (E) of different mouse genotypes. **F.** Graphs for inter-crossover distances for WT vs. *Sumo1^Gt/Gt^* (all chromosomes; left), WT vs. *Sumo1^Gt/Gt^* (matched chromosome lengths to minimize edge effects; middle), and WT vs. *Senp1^M1/M1^*(all chromosomes; right). In the middle panel, the chromosomes with matched lengths (7-12 µm) represent 48% of wild-type, n = 505; and 55% of *Sumo1^Gt/Gt^*, n = 417; median lengths = 9.47 and 9.44 um respectively, Wilcoxon rank sum test for differences in median length *p* = 0.76. PDF, Probability Density Function. **G.** Comparison of crossover interference in pachytene spermatocytes among wild type, *Sumo1^Gt/Gt^*, and *Senp1^M1/M1^* mice. CoC, Coefficient of coincidence. **H.** Representative images of metaphase I chromosome spread showing a homologous chromosome with 1, 2, or 3 chiasmata. Yellow arrows indicate the chiasma position. **I.** Representative metaphase I chromosome spreads of spermatocytes and oocytes from wild type, *Sumo1^Gt/Gt^*, and *Senp1^M1/M1^* mice, respectively. White numbers indicate chiasma counts per cell (19 autosomes in spermatocytes and all 20 chromosomes in oocytes). Scale bars are 10 μm. **J, L**. Quantification of chiasma counts per metaphase I spermatocytes (J) or oocytes (L) from different mouse genotypes. **K, M**. Fraction of individual homologous chromosomes possessing 0, 1, 2, or ≥3 chiasmata, from metaphase I spermatocytes (K) or oocytes (M) of wild type, *Sumo1^Gt/Gt^*, and *Senp1^M1/M1^*mice, respectively. Data are presented as mean ± SD (B, D, J, L). Data were analyzed with Brown-Forsythe and Welch ANOVA followed by Dunnett’s T3 multiple comparisons (B, D), ordinary one-way ANOVA followed by Dunnett’s multiple comparisons (J), Kruskal-Wallis test followed by Dunn’s multiple comparisons (L), and Fisher’s exact test (C, E, K, M). Numbers in parenthesis indicate number of nuclei (B, D, J, L) or SCs/homologs (C, E, K, M) analyzed. ns, not significant; \*\**p* < 0.01; \*\*\**p* < 0.001; \*\*\*\**p* < 0.0001.

In contrast to the *Sumo1^Gt/Gt^* mutant, *Senp1^M1/M1^*meiocytes had higher levels of crossover-specific MLH1 foci, with increases of 24.7% in spermatocytes (27.8 ± 2.3 compared to 22.3 ± 1.5 in wild type, mean ± S.D., *p*<0.0001; **Figure 4B**) and 22.1% in oocytes (31.5 ± 3.1 versus 25.8 ± 1.8 in wild type, *p*<0.0001; **Figure 4D**). *Senp1^M1/M1^*meiocytes had fewer apparent non-exchange chromosomes, with zero MLH1 foci, and higher levels of SCs with ≥2 MLH1 foci relative to wild type (*p*<0.0001 for both spermatocytes and oocytes, Fisher’s exact test, **Figure 4C** and **4E**). In *Senp1^M1/M1^*spermatocytes, just 0.4% of SCs had zero MLH1 foci compared to 2.9% in wild-type cells; while SCs with two MLH1 foci increased from 22.0% to 48.9%, with a corresponding decrease in SCs with a single MLH1 focus (**Figure 4C**). Similar trends were seen for *Senp1^M1/M1^* oocytes, with a striking increase in SCs with three MLH1 foci, which were rare in wild type (10.8% versus 4.4% in *Senp1^M1/M1^* and wild type, respectively; **Figure 4E**).

Crossover interference describes the observation that adjacent crossovers are more widely and evenly spaced than predicted for a random distribution (4, 30). Physical distance along prophase-I chromosomes is the metric for crossover interference. Thus, modulation of chromosome length *per se* could account for the changes in crossover rates observed in *Sumo1^Gt/Gt^* and *Senp1^M1/M1^*meiocytes. Alternatively, or in addition, the distance over which crossover interference spreads could be altered, with longer or shorter interference distances resulting in fewer or more crossovers in *Sumo1^Gt/Gt^* and *Senp1^M1/M1^* meiocytes, respectively. To distinguish these possibilities, crossover interference was assessed by comparing: (i) distances between adjacent crossover sites (MLH1 foci) (**Figure 4F**); and (ii) coefficient of coincidence curves (CoC curves), which plot the ratios of observed frequencies of double crossovers divided by those expected if crossovers occurred independently in pairs of intervals separated by increasing distance (**Figure 4G**)(29). CoC values of less than 1 indicate positive interference (double crossovers less likely than random expectations), a value of 1 indicates no interference, and values greater than 1 imply negative interference.

Median inter-crossover distances for all chromosomes skewed smaller in *Sumo1^Gt/Gt^* but not in *Senp1^M1/M1^* mutant spermatocytes (7.0 and 8.3 µm, respectively versus 8.4 µm in wild type; *p* <<0.01 and 0.55, Mann-Whitney U test). Taken at face value, this suggests that the *Sumo1^Gt/Gt^* mutation may weaken interference. However, the altered distribution of chromosome lengths in *Sumo1^Gt/Gt^*cells could reduce average inter-crossover distance in *Sumo1^Gt/Gt^*cells without altering interference *per se*. Specifically, the range of observable distances between adjacent crossovers will be reduced for chromosomes whose lengths approach the minimal interference distance (or multiples thereof)(31–33). Consistent with this possibility, when bins of chromosomes with comparable length distributions were compared (7-12 µm **Figure 4F**), no difference in the median inter-crossover distance was observed (6.4 µm for both *Sumo1^Gt/Gt^* and wild type, respectively, *p* = 0.58 Mann-Whitney U test). Moreover, CoC curves were comparable for crossovers in wild-type, *Sumo1^Gt/Gt^*, and *Senp1^M1/M1^*spermatocytes, with inter-interval distances of 5.5, 5.6, and 5.2 µm, respectively at a CoC of 0.5 (blue dashed line in **Figure 4G**). Thus, crossover interference appears to be largely unperturbed in *Sumo1^Gt/Gt^* and *Senp1^M1/M1^* mutants.

To confirm that crossover numbers are modulated by SUMO1 levels, chiasmata (the cytological manifestations of crossing over in condensed metaphase-I chromosomes) were counted (**Figure 4H–M**). This analysis confirmed the results from counts of MLH1 foci, with fewer chiasmata in *Sumo1^Gt/Gt^*nuclei and more chiasmata in *Senp1^M1/M1^* nuclei (**Figure 4J** for spermatocytes and **Figure 4L** for oocytes). Similarly, distributions of chiasmata per bivalent mirrored those of MLH1 foci (**Figure 4K** and **4M**), with one important exception – no univalent chromosomes, lacking chiasmata, were observed in *Sumo1^Gt/Gt^* nuclei even though 8.2% of SCs in *Sumo1^Gt/Gt^* spermatocytes and 3.5% of SCs in *Sumo1^Gt/Gt^* oocytes lacked MLH1 foci (**Figure 4C** and **4E**, respectively). The reason for this discrepancy is unclear but indicates that *Sumo1^Gt/Gt^*meiocytes remain proficient for crossover assurance, i.e., the formation of at least one crossover between every homolog pair, as required for accurate segregation at the first meiotic division.

We also tested whether the elevated crossover levels in *Senp1^M1/M1^*meiocytes were SUMO1 dependent by counting MLH1 foci in pachytene spermatocytes of *Sumo1^Gt/Gt^ Senp1^M1/M1^* double mutants (**Supplemental Figure S3A,B**). Analogous to the effects on chromosome length, the increased crossing over was largely though not completely dependent on the presence of SUMO1: MLH1 foci in *Sumo1^Gt/Gt^ Senp1^M1/M1^* nuclei averaged 22.2 ± 1.6, significantly lower than in the *Senp1^M1/M1^*single mutant (28.0 ± 3.2, mean ± S.D.; *p*<0.0001), but still higher than the *Sumo1^Gt/Gt^* single-mutant (20.2 ± 3.4, *p*=0.0022) and not different to wild-type MLH1 focus counts (22.3 ± 1.5, *p*>0.9999).

Finally, we asked whether the *Senp1^M1^* mutation had a dosage effect on crossover rate by quantifying MLH1 foci in spermatocytes of *Senp1^M1/+^* heterozygotes (**Supplemental Figure S5A**). Indeed, *Senp1^M1/+^* heterozygosity caused an intermediate effect on crossover levels, with 24.6 ± 2.0 MLH1 foci per nucleus, significantly lower than the *Senp1^M1/M1^* homozygote (27.8 ± 2.3, mean ± S.D., *p*<0.0001), but still higher than wild-type crossover levels (22.3 ± 1.5, *p*<0.0001; this result was confirmed by analysis of a second crossover-specific marker, HEI10; **Supplemental Figure S5B**).

### Early and Intermediate Steps of Recombination Are Modulated by SUMO

To determine whether earlier stages of meiotic recombination are affected by SUMO, spermatocyte chromosome spreads were analyzed for markers of DSB formation (DMC1) and intermediate steps of recombination (MSH4 and RNF212; **Figure 5**). The meiosis-specific RecA-homolog DMC1 assembles onto resected DSB ends and catalyzes homologous pairing and DNA strand exchange (34). Numbers of DMC1 foci in leptotene-stage nuclei were decreased in *Sumo1^Gt/Gt^* nuclei (227.8 ± 26.7 foci per nucleus versus 246.6 ± 30.9 in wild type, mean ± S.D., *p*=0.0243). This ∼8% reduction in DMC1 matches the ∼9% reduction in axis length seen in *Sumo1^Gt/Gt^* spermatocytes (**Figure 3B**). In contrast, DMC1 foci were increased by ∼22% in *Senp1^M1/M1^* nuclei (301.5 ± 47.3 foci per nucleus versus 246.6 ± 30.9 in wild type, mean ± S.D., *p*<0.0001; **Figure 5A, B**), a larger increase than expected from the ∼13.2% increase in axis length (**Figure 3C**).

**Figure 5.**
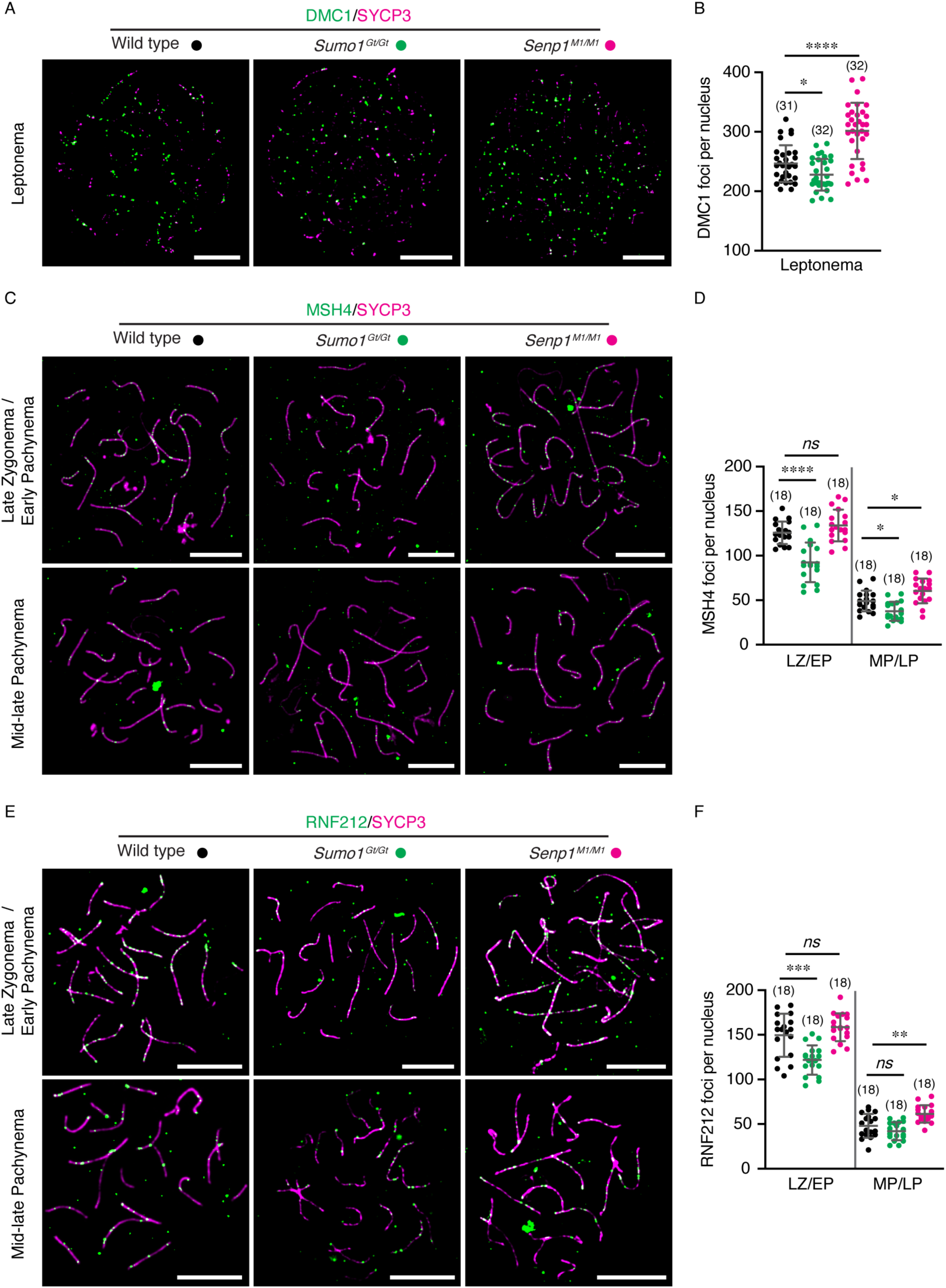
Recombination markers of early and intermediate steps are altered by SUMO1 levels. **A.** Representative surface spread images of leptotene spermatocytes immunostained for SYCP3 (magenta) and DMC1 (green) from wild type, *Sumo1^Gt/Gt^*, and *Senp1^M1/M1^* mice, respectively. **B.** Comparison of DMC1 foci counts per nucleus of leptotene spermatocytes. **C, E**. Representative images of late zygotene/early pachytene and mid-late pachytene spermatocytes immunostained for MSH4 (C) or RNF212 (E) from wild type, *Sumo1^Gt/Gt^*, and *Senp1^M1/M1^* mice, respectively. **D, F**. Quantification of MSH4 (D) or RNF212 (F) foci per nucleus of different stage of spermatocytes from wild type, *Sumo1^Gt/Gt^*, and *Senp1^M1/M1^*mice, respectively. Scale bars are 10 μm. Data are presented as mean ± SD (B, D, F). Data were analyzed with Brown-Forsythe and Welch ANOVA followed by Dunnett’s T3 multiple comparisons (B), and ordinary one-way ANOVA followed by Dunnett’s multiple comparisons (D, F). Numbers in parenthesis indicate number of nuclei examined. ns, not significant; \**p* < 0.05; \*\**p* < 0.01; \*\*\**p* < 0.001; \*\*\*\**p* < 0.0001.

The MutSγ complex, comprised of MSH4 and MSH5, binds and stabilizes nascent strand-exchange intermediates to promote homologs synapsis, and facilitates the formation and resolution of double-Holliday junctions into crossovers (35–37). Consistent with previous studies (38–40), immunostaining for MSH4 in wild-type spermatocytes revealed 125.7 ± 12.6 foci in late zygotene/early pachytene stage nuclei as cells completed synapsis, which then reduced to 49.1 ± 11.7 foci in mid/late pachytene (**Figure 5C** and **5D**). In *Sumo1^Gt/Gt^* nuclei, numbers of MSH4 foci were reduced by ∼27% in late zygotene/early pachytene (92.4 ± 22.3, ∼27%) and ∼24% mid/late pachytene nuclei (37.6 ± 10.7, mean ± S.D., *p*<0.0001 and *p*=0.0123, respectively), larger decreases than the ∼9% reduction in axis length in this mutant. In the *Senp1^M1/M1^* mutant, numbers of MSH4 foci were not significantly different from wild type in late zygotene/early pachytene nuclei (133.9 ± 17.8 versus 125.7 ± 12.6, mean ± S.D., *p*=0.2968), but were higher in mid/late pachytene (60.4 ± 13.8 versus 49.1 ± 11.7, mean ± S.D., *p*=0.0143), consistent with the higher numbers of crossover-specific MLH1 foci formed in this mutant (above and **Figure 4**).

Finally, we analyzed localization of the RING E3 ligase, RNF212, which is required to stabilize MutSγ at crossover sites (**Figure 5E**)(17, 38, 39). The dynamics of RNF212 in wild-type spermatocytes were similar to those of MSH4, with 149.5 ± 24.1 foci in late zygotene/early pachytene nuclei and 48.1 ± 13.3 foci in mid/late pachytene (**Figure 5F**, mean ± S.D.). In the *Sumo1^Gt/Gt^* mutant, RNF212 foci were significantly reduced in late zygotene/early pachytene but not in mid/late pachytene nuclei, although the trend was slightly lower (121.9 ± 16.4 and 42.1 ± 10.1, respectively; *p*=0.0001 and *p*=0.1903). In *Senp1^M1/M1^*mutant nuclei, numbers of RNF212 foci in late zygotene/early pachytene trended slightly higher but were not significantly different from wild type (158.5 ± 15.6 versus 149.5 ± 24.1, mean ± S.D., *p*=0.2758). However, RNF212 focus numbers were significantly increased during mid/late pachytene (61.4 ± 9.7 versus 48.1 ± 13.3, mean ± S.D., *p*=0.0015). Thus, SUMO1 level quantitatively modulates all steps of meiotic recombination from DSB formation through crossing over.

## DISCUSSION

Our analysis shows that: (i) axis-associated SUMO correlates with the longer homolog axes, shorter chromatin loops (11), increased DSB levels (12), and higher crossover rates seen in oocytes relative to spermatocytes (heterochiasmy); and (ii) changes in SUMOylation can modulate chromosome length and crossover rate without impacting interference. Previous studies in human and mouse indicated that the processes responsible for heterochiasmy do not act by altering the strength of interference but by altering chromosome axis length (9, 11, 29, 41, 42). However, more recent studies in mouse indicate that this model is not generalizable across strains and subspecies. Notably, heterochiasmy is reversed in some subspecies, with males having more crossovers than females, even though chromosome length remains shorter in males (15, 43). Thus, while SUMO might contribute to heterochiasmy in some cases, it seems unlikely to be the sole driver. DSB number, the efficiency of converting DSBs into crossovers, and interference are other facets of recombination that can show significant sex and strain-specific differences (15, 43).

Analysis of *Sumo1^Gt^* and *Senp1^M1^* mutants reveal that levels of SUMO1 conjugation reciprocally modulate the lengths of chromatin loops and homolog axes, with associated effects on numbers of DSBs and crossovers. We suggest that SUMO-mediated regulation of two distinct but coupled features of meiotic chromosomes account for modulation of DSB number and crossover frequency, respectively. Loop-axis modules (or multiples thereof) are inferred to be the minimum units of DSB formation, with breaks forming within protein complexes associated with the bases of the loops (3, 44–46). Thus, numbers of potential DSB sites will increase when chromatin loops are smaller and more numerous, as seen in oocytes (12) and in *Senp1^M1/M1^*mutant meiocytes; and *vice versa* for spermatocytes and *Sumo1^Gt/Gt^*meiocytes. Physical distance along prophase-I homolog axes is the metric for crossover interference (4). Thus, increased axis length per se will proportionally increase crossover number even if the strength of crossover interference remains unchanged, as seen in *Senp1^M1/M1^*and *Sumo1^Gt/Gt^* mutants.

The higher incidence of SCs that lack an MLH1 focus in the *Sumo1^Gt/Gt^*mutant focus raises the possibility that crossover assurance may be compromised in the absence of SUMO1 (**Figure 3**). However, univalents were not observed in metaphase-I nuclei suggesting that crossover assurance remains efficient. This disparity could be explained if occasional failure of the major MLH1-dependent pathway can be compensated by an alternative crossover resolution enzyme. A likely candidate would be the structure-selective endonuclease, MUS81-EME1 (43, 47). Alternatively, SCs lacking MLH1 foci in *Sumo1^Gt/Gt^* cells might reflect asynchronous assembly/disassembly and/or shorter lifespans of MLH1-marked crossover complexes across chromosomes in the absence of SUMO1.

### How could SUMO modulate axis length?

The distinct oligomerization and interaction properties of SUMO1 and SUMO2/3 help determine both their target preferences and the consequences of modification (48, 49). SUMO2/3 can assemble into chains linked via lysines in their disordered N-termini. By contrast, SUMO1 is generally assumed to modify proteins as a monomer or to cap poly-SUMO2/3 chains to terminate their oligomerization. Non-covalent SUMO binding by SUMO-interaction motifs (SIMs) and other motifs (e.g., WD40 repeats (50)) can promote interactions with partner proteins. Moreover, paralog- and oligomer-selective SIMs can provide additional levels of specificity (51). SUMO is also broadly implicated in protein quality control with both binding and conjugation of SUMO1 conferring a chaperone-like function to enhance protein solubility, whereas conjugation of poly-SUMO2/3 chains can promote protein degradation in collaboration with the ubiquitin-proteasome system (48, 49). However, these distinctions are not absolute. For example, mono-SUMO2 (or short chain) modification mediates protein interactions (51, 52) and likely improves protein solubility (53); and SUMO1-capped poly-SUMO2/3 chains are the preferred substrate of the SUMO-targeted ubiquitin ligase RNF111 (21).

Thus, the shorter axes associated with loss of SUMO1 modification in *Sumo1^Gt/Gt^* meiocytes could result from decreased solubility of pertinent targets, altered protein interactions, and/or increased turnover due to conjugation of poly-SUMO2/3 chains (the growth of which are not attenuated by SUMO1 capping). On the other hand, the longer axes of *Senp1^M1/M1^* meiocytes could reflect enhanced solubility and/or stability of SUMO1 conjugates, altered protein interactions, and/or stabilization of SUMO2/3 conjugates because excess SUMO1 limits chain growth.

Analysis of the *Senp1^M1/M1^ Sumo1^Gt/Gt^* double mutant suggests that SENP1 influences chromosome length and crossover number through SUMO1-dependent and independent pathways. Sharma et al. provided evidence that SENP1 can indirectly modulate SUMO2/3 conjugation by influencing the stability of a second isopeptidase, SENP2 (20). SENP1 is also suggested to regulate JAK/STAT signaling through direct interactions with JAK2 and STAT3 (54, 55). Intriguingly, non-canonical JAK2 signaling influences expression of the cohesin component STAG3, which has the potential to alter axis length (56).

### SUMO targets that might modulate axis length

Given the thousands of known SUMO targets (48, 57, 58), effects of *Sumo1^Gt/Gt^* and *Senp1^M1/M1^* mutations will be pleiotropic; and direct and/or indirect effects could modulate axis length. Components of homolog axes that are known to modulate their lengths are logical SUMO targets and include cohesin, condensin, and meiosis-specific axis components (3, 26–28, 59–64). All these components are SUMOylated during meiosis in both budding yeast (57) and likely in mammals (58, 65).

Cohesin is an attractive candidate for SUMO-modulated regulation of axis length. In somatic cells, SUMO mediates the establishment of cohesion, and the stabilization and turnover of cohesin complexes (66–71). Initial mono-SUMOylation may facilitate establishment of cohesion while subsequent multi- or poly-SUMOylation triggers dissociation or degradation of cohesin. Notably, in somatic cells, the isopeptidase SENP6 limits the polySUMOylation and turnover of cohesin from chromatin though an interaction with the cohesin regulator PDS5 (71), defining a pathway that appears to be conserved in various fungi (70, 72, 73). In somatic cells, human PDS5A and PDS5B regulate the number and size of cohesin-mediated chromatin loops (74, 75); and co-depletion of PDS5A and PDS5B in mouse spermatocytes results in shorter homolog axes (64). During meiosis in Sordaria, Spo76/Pds5 protects cohesin from proteasomal degradation mediated by the SUMO-targeted ubiquitin ligase Slx8 (73). However, while the shorter meiotic chromosomes seen in budding yeast *pds5* mutants are dependent on the ubiquitin-proteasome system, SUMOylation is not involved (61). Ubiquitylation mediated by the E2 enzymes Rad6/HR6A/B also intersects with SUMO (76, 77) and *Hr6b* knock-out mice have longer chromosomes and increased crossing over (78). Finally, SUMO-mediated regulation of S-phase (79–83) could modulate cohesin loading at replication forks (84).

Condensin subunits are SUMOylated during meiosis (57, 58) and condensin mutants have longer prophase-I chromosomes and more crossovers (85–88). Moreover, condensin turnover also appears to be regulated by SUMO (71, 72, 89). Meiosis-specific homolog axes proteins, including mammalian SYCP3 and HORMAD1, are also SUMOylated (57, 58, 65), implicated in protecting cohesins (73), and mouse *Sycp3* mutants have longer homolog axes (28).

General components of chromatin, including histones, remodelers and transcription factors, are also prominent meiotic targets of SUMO (57), which can alter chromatin organization and axis length (90), and might do so in ways that might reflect sex-specific epigenetic states (91, 92).

### Could stress-modulated SUMOylation contribute to crossover-rate dimorphism, plasticity, and evolvability?

Various types of cellular stress cause global changes in both the level and pattern of proteome SUMOylation (93, 94). Consequently, systemic, tissue-specific, or cellular stress levels may cause global SUMOylation to vary among individuals, tissues, or cells. Such variation could contribute the observed differences in chromosome axis lengths and crossover rates between males and females, individuals of the same sex, and individual meiocytes (6, 13). Moreover, per-cell variation in global SUMOylation provides an attractive model to explain the covariation of axis lengths and crossover rates across different chromosomes within individual meiocytes; and the overdispersion of hyper- and hypo-crossover cells (6). Stress-induced SUMOylation may also contribute to the phenomenon of stress-induced recombination (95). Both stress-induced recombination and per-cell variation in crossover rate are inferred to enhance evolvability (6, 95, 96). Thus, the SUMO-stress response could contribute to evolutionary adaptation by translating environmental variation to changes in the rate of meiotic recombination.

## Acknowledgement

We thank Nancy Kleckner, Richard Shultz, Satoshi Namekawa and members of the Hunter lab for helpful discussions. This work was supported in part by award R01HD109322 to N.H. from the Eunice Kennedy National Institute of Child Health and Development. Y.Y. is supported by a Guangdong Basic and Applied Basic Research Foundation (2024A1515012907), and H.B.D.P.R is supported by DBT-Ramalingaswami re-entry fellowship and NIAB core grant C0031. N.H. is an Investigator with the Howard Hughes Medical Institute.

## Author contributions

Y.Y., H.B.D.P.R and N.H. conceived the study and designed the experiments. H.B.D.P.R, H.Q., S.S., W.Q., S.B., A.D., S.B., A.S., L.B., H.T., and B.V. performed the experiments and analyzed the data. Y.Y. performed oocyte culture experiments and prepared figures. M.W. performed crossover interference analysis. Y.Y., H.B.D.P.R and N.H. wrote the manuscripts with input from M.W. All of authors edited the manuscript.

## Competing financial interests

The authors declare no competing financial interests.

## MATERIALS AND METHODS

### Animals

*Sumo1^Gt(XA024)Byg^* and *Senp1^M1Mku^* mutant lines were previously described (18–20) and were made congenic with the C57BL/6J background. Animals were housed up to four per cage in a local vivarium with temperature control and a 12-hour light/dark cycle, and *ad libitum* access to standard rodent chow and water. Mice were maintained and used for experimentation according to the guidelines of the Institutional Animal Care and Use Committees of the University of California, Davis.

### Surface Spreads of Prophase-I Spermatocyte and Oocyte Chromosomes

Testes were dissected from freshly euthanised adult animals (2-6-month-old) and processed for chromosome spreads as described (47, 97). Briefly, testes were decapsulated and seminiferous tubules incubated in freshly made hypotonic extraction buffer (HEB; 50 mM sucrose, 30 mM Tris–HCl pH 8.0, 17 mM trisodium citrate, 5 mM EDTA, 0.5 mM DDT, 0.5 mM PMSF) on ice for 20 min. Tubules were minced on glass depression slides with a small amount of 0.1 M sucrose (Sigma-Aldrich, S0389), spread on glass slides coated in 1% paraformaldehyde with 0.15% Triton X-100 pH 9.2, and air-dried overnight in a humid chamber. Fetal ovaries were dissected from E16.5-18.5 female fetuses and processed for surface spreading of prophase-I oocyte chromosomes as described (97). Ovaries were also incubated in HEB on ice for 20 mins and then minced in a 20 µl drop of 0.1 M sucrose. The cell suspension was spread on a glass slide coated in 1% paraformaldehyde with 0.15% Triton X-100 pH 9.2 and air-dried overnight in a humid chamber. Finally, the slides were washed in 0.4% Photo Flo (Kodak, 1464510) and dried again at room temperature for immunostaining analysis.

### Chromosome Spreads of Metaphase-I Spermatocytes

For quantification of chiasmata, chromosome spread from diakinesis/metaphase I stage spermatocytes were prepared using the technique described (97). Briefly, Adult mouse testes were dissected and decapsulated into hypotonic solution (1% Sodium Citrate). Seminiferous tubules were minced to make a cell suspension. Following centrifugation, the cell pellet was slowly suspended in methanol/acetic acid/chloroform (3:1:0.015). Cells were harvested by centrifugation, and the pellet was slowly re-suspended in ice-cold methanol/acetic acid (3:1). Aliquots of the cell suspension were dropped onto clean glass slides and air dried at room temperature. For imaging, DNA was stained with DAPI.

### Oocyte Culture and Metaphase-I Chromosome Spreads

Germinal-vesicle stage oocytes were retrieved from freshly dissected ovaries of 21-25-day old animals without prior hormonal stimulation. Only oocytes with integral cumulus cell layers were utilized. Oocytes were collected in M2 medium (Sigma-Aldrich, M7167) under mineral oil (Sigma-Aldrich, M8410 and Nidacon, NO-100) at 37°C and cultured for *in vitro* maturation after mechanically removing surrounding cumulus cells for observation of germinal vesicle breakdown (GVBD). To count chiasmata, metaphase-I oocytes were acquired after 7 hrs of culture, and then prepared for chromosome spreads as previously described with modifications (98). Briefly, Acid Tyrode’s solution (Sigma-Aldrich, M1788) was applied to remove zona pellucida, and zona-free oocytes were incubated in hypotonic solution (1% Sodium Citrate) for 20 min, followed by fixation in fixing solution (methanol: acetic acid = 3:1) for 5 min. The cell suspension was dropped onto a clean microscope slide, and spreads were air dried at room temperature. DNA was stained with DAPI for imaging.

### Immunofluorescence Cytology and Image Acquisition

Immunofluorescence staining was performed as previously described (97). For SUMO1, SUMO2/3, and HEI10 staining, extra pepsin and DNase treatments were applied prior to the blocking procedure. Specifically, slides were incubated with 1 mg/ml pepsin in 0.01 N HCl for 3 minutes at room temperature, followed by 0.01 mg/ml DNase treatment for 15 minutes at 37 °C. After blocking slides at room temperature for 1 hour, the following primary antibodies were used for incubation overnight at room temperature: mouse anti-SYCP3 (Santa Cruz, sc-74568; 1:200), rabbit anti-SYCP3 (Santa Cruz, sc-33195; 1:200), human anti-centromere antibodies (ACA or CREST; ImmunoVision, HCT-0100; 1:1000), mouse anti-SUMO1 (SUMO-1 21C7; 1:150), mouse anti-SUMO2/3 (SUMO-2 8A2; 1:150), mouse anti-MLH1 (BD Pharmingen, 550838; 1:50), rabbit anti-DMC1 (Santa Cruz, sc-22768; 1:200), rabbit anti-MSH4 (Abcam, ab58666; 1:100), mouse anti-CCNB1IP1/HEI10 (Abcam, ab118999; 1:150), guinea pig anti-RNF212 (1:50)(38). After washing, slides were subsequently incubated with the following goat secondary antibodies for 1 hr at 37 °C: anti-mouse 488 (Thermo Fisher Scientific, A11029; 1:1000), anti-mouse 594 (Thermo Fisher Scientific, A11020; 1:1000), anti-rabbit 488 (Thermo Fisher Scientific, A11070; 1:1000), anti-rabbit 568 (Thermo Fisher Scientific, A11036; 1:1000), anti-human DyLight 649 (Jackson Labs, 109-495-088; 1:200), anti-guinea pig fluorescein isothiocyanate (FITC; Jackson Labs, 106-096-006; 1:200). Coverslips were mounted with ProLong Diamond antifade reagent (Thermo Fisher Scientific, P36970) prior to imaging.

All images were acquired using a Zeiss AxioPlan II microscope equipped with a 63x Plan Apochromat 1.4 NA objective and EXFO X-Cite metal halide light source. Images were captured by a Hamamatsu ORCA-ER CCD camera and Image processing, and analysis were performed using Volocity (Perkin Elmer) and ImageJ (NIH) software.

### FISH on Surface Spreads of Prophase-I Spermatocyte Chromosome

FISH staining was performed as described (25). All staining processes were performed in the dark to preserve fluorescence signal. Briefly, following immunofluorescence staining, slides were fixed with 2% paraformaldehyde in 1 X PBS for 10 min at room temperature, and then washed in PBS. The slides were then sequentially dehydrated with 70% ethanol for 4 min, 90% ethanol for 4 min and 100% ethanol for 5 min, and air-dried. Mouse chromosome 5 probe (XMP 5 green; MetaSystems Probes, D-1405-050-F1) in hybridization solution was applied, and DNA was denatured by placing slides on a heating block for 7 min at 79 °C. The probe was then hybridized overnight by incubating slides at 37 °C. Slides were washed in 0.1X SSC, 0.4X SSC with 0.3% NP-40, 1X PBS with 0.05% Tween-20, and H_2_O, respectively, followed by mounting with ProLong Diamond antifade reagent for imaging.

### Quantification and Statistical Analysis

For SUMO1 and SUMO2/3 staining, the large staining signals associated with the sex chromosomes and at centromeres were not included in quantifications. To quantify the localization of MLH1, SUMO1, SUMO2/3, DMC1, MSH4, RNF212 and HEI10, immunostaining, foci closely juxtaposed to the SYCP3-staining chromosome axes were counted. Coefficient of coincidence analysis was performed using the MATLAB function getCoC, as previously described (99). Chromosomes were divided into 20 intervals, corresponding to an average interval size of 0.5 – 0.6 um.

Contingency table data were analyzed using Fisher’s exact tests. Data are presented as mean ± SD. For comparisons between two groups, an unpaired t-test (parametric) or Mann–Whitney U test (nonparametric) was used. For multiple comparisons, parametric data with equal variances were analyzed using one-way ANOVA followed by Dunnett’s post hoc test. When variances were unequal, Brown–Forsythe and Welch ANOVA were performed, followed by Dunnett’s T3 post hoc test. Nonparametric data were analyzed using the Kruskal–Wallis test followed by Dunn’s post hoc test. Data were processed using GraphPad Prism 10, with a significance threshold of p < 0.05. In all comparisons, at least three age-matched mice were examined from at least two independent experiments.

## Notes

### Competing Interest Statement

The authors have declared no competing interest.

