## Supplemental Figures for "SUMO mediates the coordinate regulation of meiotic chromosome length and crossover rate"

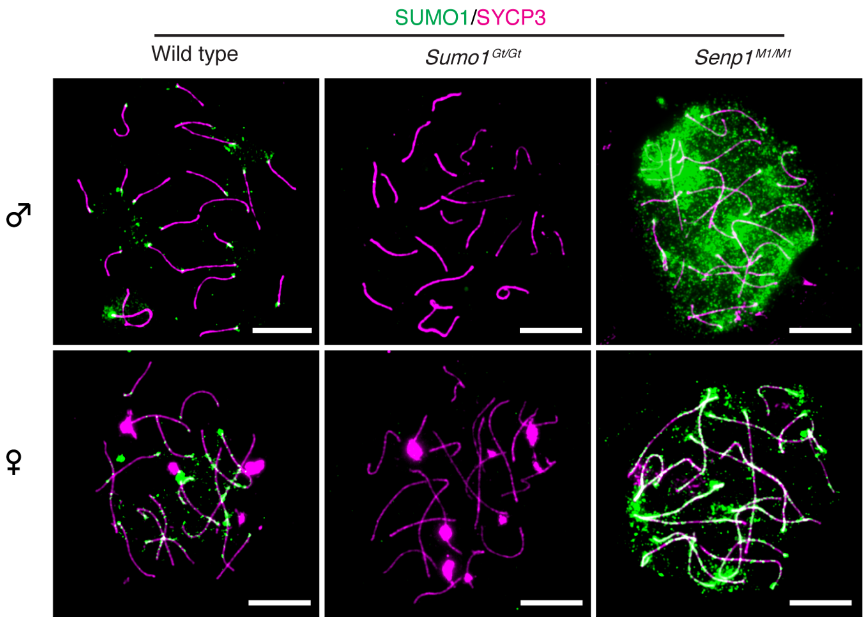


**Figure S1. SUMO1 levels in pachytene spermatocytes and oocytes of gene-modified mouse models.**

Representative surface spread images of pachytene spermatocytes and oocytes immunostained for SYCP3 (magenta) and SUMO1 (green) from wild type, *Sumo1*$^{Gt/Gt}$, and *Senp1*$^{M1/M1}$ mice, respectively. Scale bars are 10 μm.

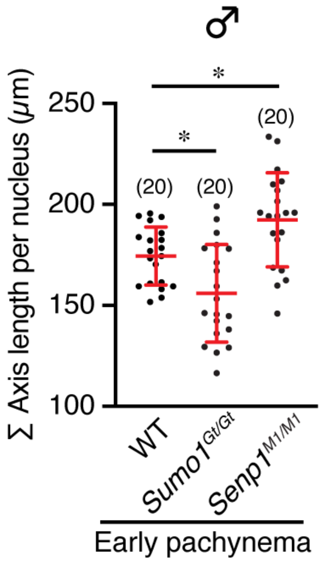


**Figure S2. Total chromosome axis length comparison in early pachytene spermatocytes of gene-modified mouse models.**

Data are presented as mean ± SD, and were analyzed with ordinary one-way ANOVA followed by Dunnett's multiple comparisons. Numbers in parenthesis indicate number of nuclei analyzed. $^*p < 0.05$.

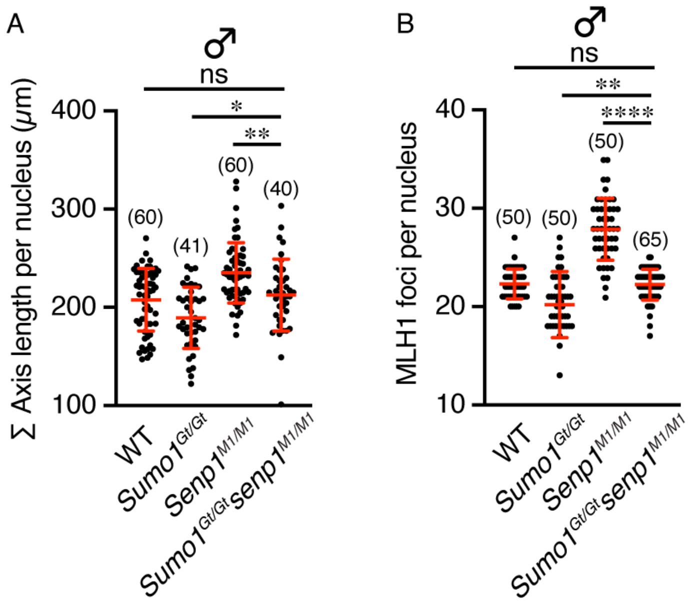


**Figure S3. Total chromosome axis length and MLH1 levels in mid-late pachytene spermatocytes of different gene-modified mouse models.**

**A, B**. Quantification of total axis length (A) or MLH1 foci (B) per nucleus of mid-late pachytene spermatocyte (19 autosomes) from wild type, *Sumo1*$^{Gt/Gt}$, *Senp1*$^{M1/M1}$, and *Sumo1*$^{Gt/Gt}$ *Senp1*$^{M1/M1}$ double mutant mice, respectively.

Data are presented as mean ± SD and were analyzed with Kruskal-Wallis test followed by Dunn's multiple comparisons. Numbers in parenthesis indicate number of nuclei analyzed. ns, not significant; *$p < 0.05$; **$p < 0.01$; ****$p < 0.0001$.

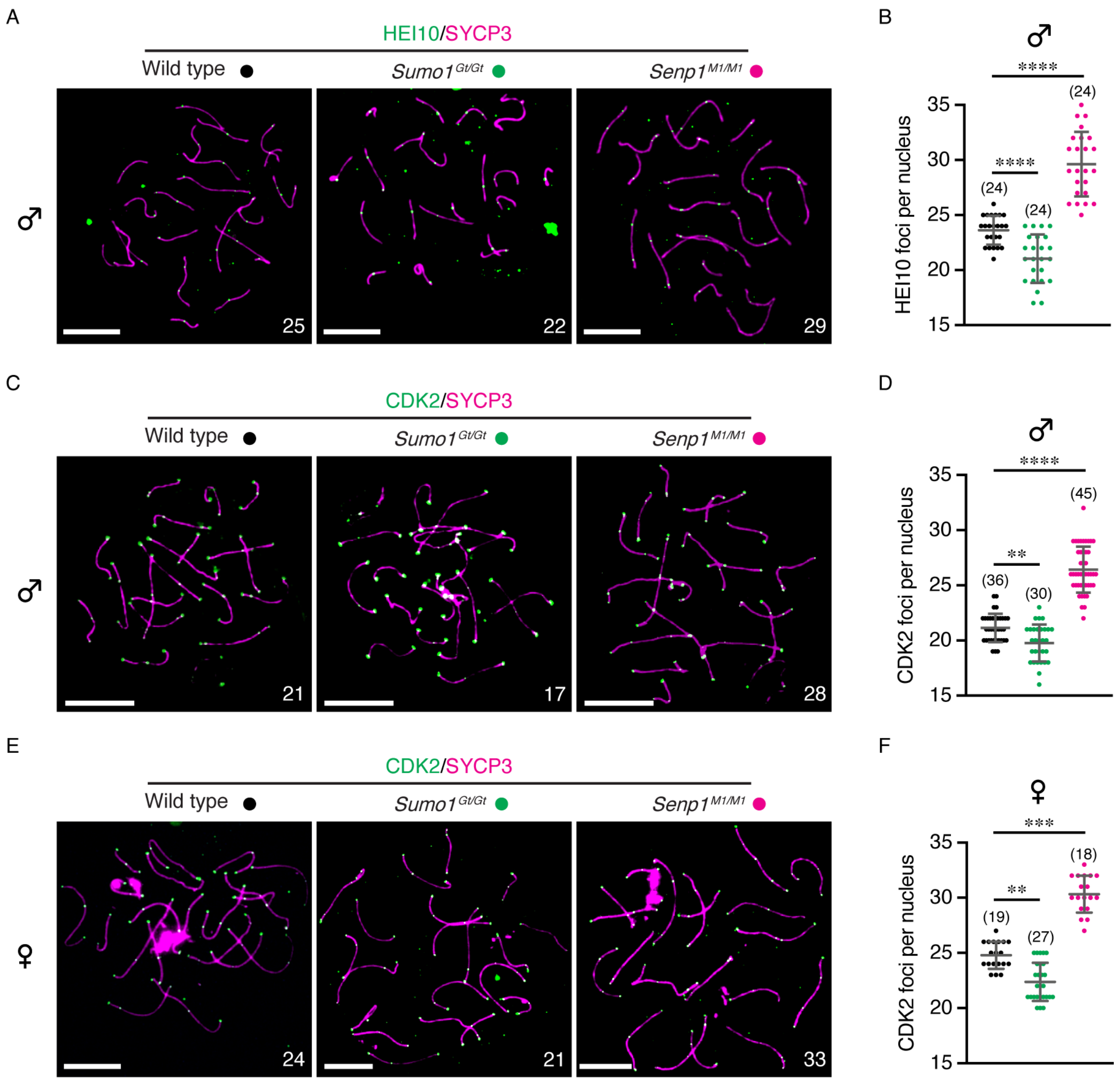


**Figure S4. Recombination markers HEI10 and CDK2 in both spermatocytes and oocytes are altered by SUMO1 levels.**

**A, C**. Representative surface spread images of mid-late pachytene spermatocytes immunostained for HEI10 (A) or CDK2 (C) from wild type, *Sumo1*$^{Gt/Gt}$, and *Senp1*$^{M1/M1}$ mice, respectively.

**B, D**. Quantification of HEI10 (B) or CDK2 (D) foci per nucleus of mid-late pachytene spermatocytes (19 autosomes) from wild type, *Sumo1*$^{Gt/Gt\,-}$, and *Senp1*$^{M1/M1}$ mice, respectively.

**E**. Representative images of pachytene oocyte surface spread immunostained for CDK2 from wild type, *Sumo1*$^{Gt/Gt}$, and *Senp1*$^{M1/M1}$ mice, respectively.

**F**. Quantification of CDK2 foci per nucleus of pachytene oocyte as in E.

White numbers indicate HEI10 (A) or CDK2 (C, E; telomeric CDK2 foci were excluded) foci counts per nucleus. Only 19 autosomes were analyzed for spermatocytes (A-D) and all 20 chromosomes were counted for oocytes (E, F). Scale bars are 10 μm (A, C, E). Data are presented as mean ± SD (B, D, F), and were analyzed with Brown-Forsythe and Welch ANOVA followed by Dunnett's T3 multiple comparisons (B), ordinary one-way ANOVA followed by Dunnett's multiple comparisons (D), and Kruskal-Wallis test followed by Dunn's multiple comparisons (F). Numbers in parenthesis indicate number of spermatocyte (B, D) or oocyte (F) nuclei analyzed. **$p < 0.01$; ***$p < 0.001$; ****$p < 0.0001$.

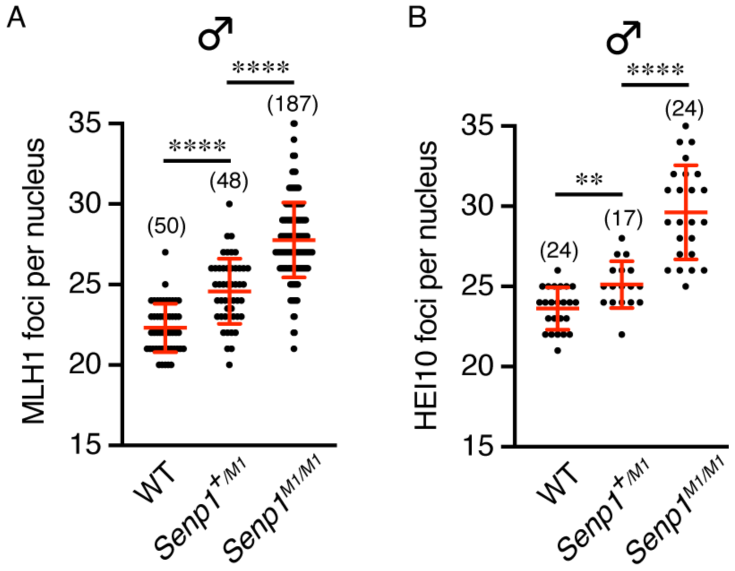


**Figure S5. Crossover levels in mouse spermatocytes with different *Senp1* genotypes.**

**A, B**. Quantification of MLH1 (A) or HEI10 (B) foci counts per nucleus of mid-late pachytene spermatocytes from wild type, *Senp1*$^{+/M1}$, and *Senp1*$^{M1/M1}$ mice, respectively.

Data are presented as mean ± SD, and were analyzed with Brown-Forsythe and Welch ANOVA followed by Dunnett's T3 multiple comparisons. Numbers in parenthesis indicate number of nuclei examined. **$p < 0.01$; ****$p < 0.0001$.
